# Actions obey Weber’s law when precision is evaluated at goal attainment

**DOI:** 10.1101/2024.07.15.603509

**Authors:** Ivan Camponogara, Robert Volcic

**Author notes:** Correspondence concerning this article should be addressed to Robert Volcic, New York University Abu Dhabi, P.O. Box 129188, Abu Dhabi, UAE.

## Abstract

Weber’s law, the psychophysical principle stating that the just noticeable difference (JND), which is inversely related to sensory precision, increases proportionally with the magnitude of a stimulus, impacts all domains of human perception, but it is violated in visually guided grasping actions. The underlying reasons for this dissociation between perception and action are still debated, and various hypotheses have been put forward, including a different coding of visual size information for perception and action, the use of positional information to guide grasping, biomechanical factors, or sensorimotor calibration. To contrast these hypotheses, we investigated Weber’s law in a new action task, the two-finger pointing task. Participants reached and touched two targets positioned at different distances apart by using their index finger and thumb. Consistent with Weber’s law, we found that the standard deviation (SD) of the final inter-finger separation, serving as a measure analogous to the JND, increased with larger inter-target distances. These findings suggest that, when considering measures strongly related with the attainment of the action goal, Weber’s law is regularly at play.

## Introduction

Every day we collect information through our diverse sensory systems and use this information to make decisions and interact with our surroundings. However, although our sensations feel accurate and truthful, they do not necessarily reflect the physical properties of the external world. For this reason, a recurrent challenge in neuroscience is understanding the relationship between physical and perceptual worlds. A crucial attempt in this direction was the work of E. H. Weber (1795–1878), who investigated the relationship between the physical intensity of a stimulus and its internal sensory representation. Weber (1834) reported that the JND, which is inversely related to sensory precision, increases proportionally with the magnitude of a stimulus. This empirical regularity, later formalized by Fechner (1860) as “Weber’s law” can be applied to a variety of taxa, sensory modalities, and measurement procedures. For this reason, Weber’s law has been hailed as one of the most reliable principles in the domain of perception (Teghtsoonian, 1971).

However, as the adage goes, for every rule there is an exception. This is the case of the visual coding for action, which violates Weber’s law (Ganel, 2015; Ganel et al., 2008, 2014). In grasping movements, before the hand has any contact with the object, the index finger and thumb reach a maximum grip aperture (MGA), which is linearly related to object size, with larger MGAs for larger objects (Jeannerod, 1986; Smeets & Brenner, 1999). Ganel et al. (2008) found that the standard deviation of MGA (SD_MGA_), serving as a measure analogous to the JND, is independent of object’s size, in stark contrast with perceptual estimates, where the JND increases with larger objects. Ganel et al. (2008) proposed that the absence of Weber’s law in action is due to the fundamental differences in how the ventral stream and the dorsal stream process visual information, in line with the Two-Visual Systems hypothesis (TVS; Goodale & Milner, 1992; Goodale, 2014).

The ventral stream, projecting from early visual areas to the inferior temporal cortex, mediates vision-for-perception, and it is based on relative metrics, as it takes into consideration the object size relative to general aspects of the visual scene. The dorsal stream, running from early visual areas to the posterior parietal cortex, mediates vision-for-action and processes the actual size of the object by means of absolute (objective) metrics, which leads to the violation of Weber’s law. Even though many studies have corroborated the violation of Weber’s law in visually guided actions, the TVS interpretation has been challenged on multiple fronts, and other hypotheses have been put forward.

The Digits-In-Space hypothesis (DIS; Smeets & Brenner, 1999, 2008) suggests that grasping parameters, including the MGA, are based on the positional information about contact points, and no internal representation of object size is needed to control grasping movements. Weber’s law, on the other hand, can only hold for properties that have a magnitude, and it does not extend to positional information. As a result, the absence of Weber’s law in grasping is a mere consequence of the fact that grasping kinematics and Weber’s law rely on different sources of information.

An alternative explanation is provided by the Biomechanical Factors hypothesis (BF; Bruno et al., 2016; Löwenkamp et al., 2015; Schenk et al., 2017; Uccelli et al., 2021; Utz et al., 2015), according to which the violation of Weber’s law might depend on the biomechanical constraints of the hand rather than on the visual dissociation between perception and action. Utz et al. (2015) argued that, as the object grows larger, hand opening approaches the limits of the absolute hand span, which artificially reduces the amount of possible variability in MGA and masks the effects of Weber’s law.

Finally, the Sensorimotor Calibration hypothesis (SMC) posits that the differences in how perceptual and action tasks follow Weber’s law might be influenced by the accessibility to sensory feedback (Bingham & Mon-Williams, 2013; Bruno & Franz, 2009; Camponogara & Volcic, 2019, 2021a, 2021b, 2022; Cesanek et al., 2018; Schenk, 2012; Volcic & Domini, 2016, 2018). Indeed, visual and haptic feedback, which is missing during perceptual estimations, may promote on-line adjustments or motor calibration, which in turn, may counteract the effect of Weber’s law by minimizing movement variability (Davarpanah Jazi et al., 2015; Heath et al., 2019).

A potential limitation of previous research is that generally, with just a few exceptions (Derzsi & Volcic, 2023), the presence of Weber’s law in action has been examined exclusively by measuring SD_MGA_ in grasping tasks, which begs the question about how and to what extent does the violation of Weber’s law generalize to other actions. Moreover, it should be noted that, even if the violation of Weber’s law is a strong empirical regularity, so far, there is still a contention about whether or not such violation can provide valuable insights into the neural control of action (Schenk et al., 2017).

With this in mind, the overarching aim of the present study is to understand the origin of this potential perception-action dissociation, and test the prediction of each hypothesis in a novel action task. For this purpose, we used a two-finger pointing task. Participants were required to reach and touch, with the index finger and thumb, two targets placed at different distances from each other, similarly to hitting a two-note chord on a piano. As a dependent variable, we considered the SD of the final inter-finger separation, that is, the distance between the index and thumb when the targets were touched (i.e., final landing points of index and thumb).

The two-finger pointing task offers a critical advantage over the grasping task, as it allows the measurement of movement precision when the goal of the action is actually attained. In contrast, in grasping tasks, the MGA occurs well before the movement’s end, and its precision may be thus influenced by other sources of noise and not necessarily be related only to the goal of the action. Obviously, the final grip aperture cannot be used to assess movement precision, as it always matches the object’s size and thus provides no additional information.

Interestingly, a recent study (Vannuscorps et al., 2022) suggested that the engagement of vision-for-perception or vision-for-action may be additionally modulated by the type of the to-be-pointed targets. Here, we tested this additional hypothesis by implementing two sets of target stimuli consisting of two unconnected dots or two dots connected by a 2D line. The rationale of this idea is that pointing two unconnected dots might represent a more realistic movement, relying on vision-for-action. Conversely, pointing the ends of a 2D line might be more akin to pantomimed grasping, relying on vision-for-perception (Whitwell & Goodale, 2022).

## Experiment 1

In Experiment 1, participants were asked to reach out and simultaneously touch, with the index finger and thumb, two unconnected dots (Points condition) or two dots connected by a line (Line condition), in an open-loop task in which visual information was removed just after the beginning of the movement (Figure 1A). This manipulation was introduced to better separate the predictions of the different hypotheses. According to the TVS hypothesis, actions rely on the dorsal stream processing as long as vision of the targets is available during movement initiation (e.g., Goodale et al., 2004, 2005; Westwood et al., 2003). Therefore, because actions should reflect absolute object metrics, a violation of Weber’s law is expected in both conditions. Possibly, if the Points condition is based on vision-for-action and the Line condition on vision-for-perception, we expect to find Weber’s law only for the Line condition (Whitwell & Goodale, 2022). From the perspective of the DIS hypothesis, actions should evade Weber’s law in both conditions, as the two-finger pointing kinematics should be coded in relation to the individual positional information of the targets, and not in relation to their mutual distance. Instead, if Weber’s law is usually violated because of biomechanical constraints (BF hypothesis), we expect that the SD of the final inter-finger separation follows Weber’s law in both conditions, because in the present study the maximum distance between targets is considerably smaller than the maximum hand span of each subject, and consequently the biomechanical factors should play no role. Finally, according to the SMC hypothesis, Weber’s law should be again observed in both conditions, because the absence of visual information after the beginning of the movement eliminates online visual corrections and prevents visuomotor calibration by concealing the outcome of the actions. These predictions are summarized in Figure 1B, Exp. 1.

**Figure 1.**
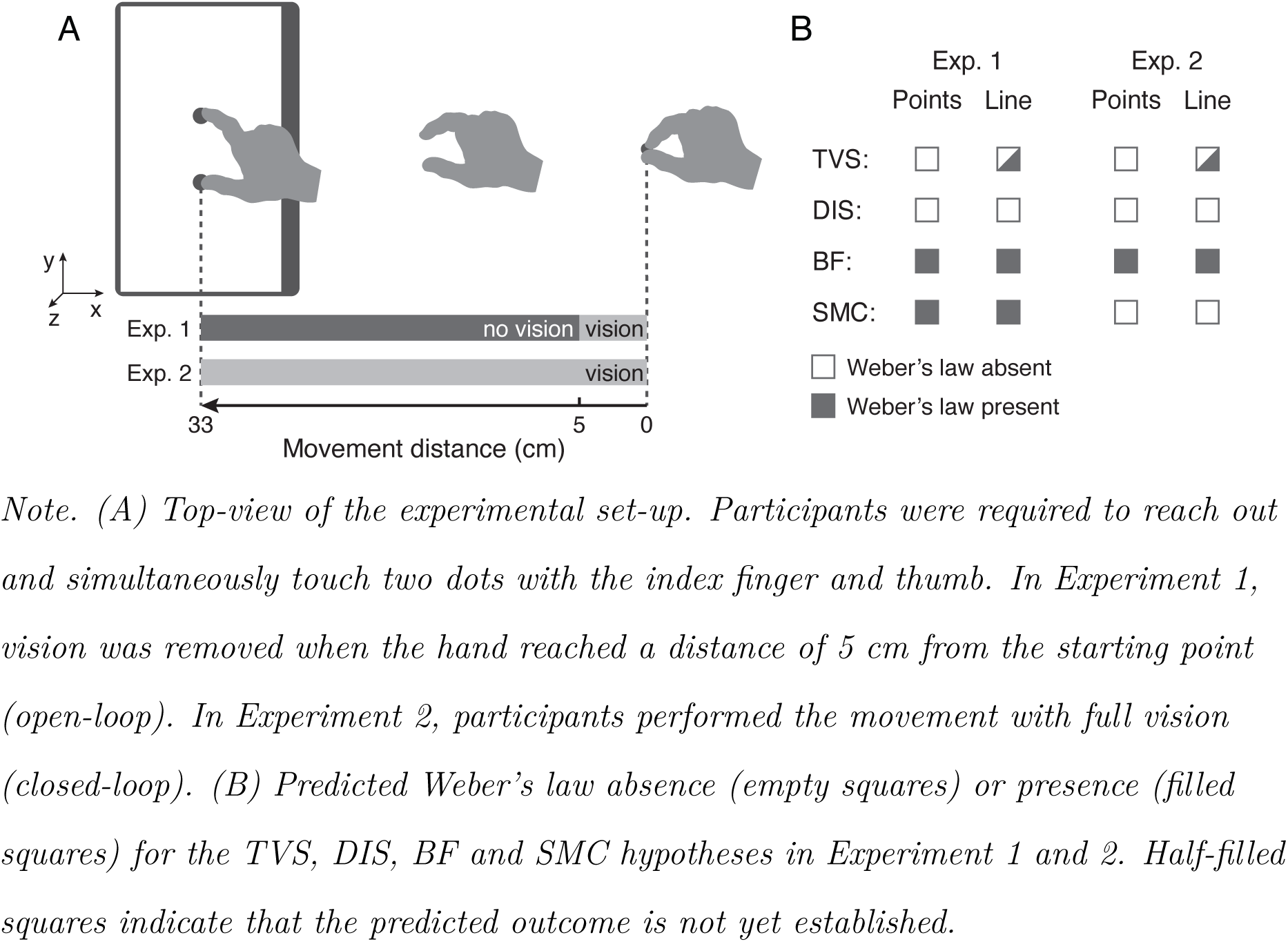
Experimental Setup and Predictions.

## Methods and Materials

### Participants

Twenty participants (mean age = 21.4 years, SD = 1.4 years, ten males) were recruited from the student population of the New York University Abu Dhabi. All participants were naïve with respect to the purpose of the study, right-handed, they had normal or corrected-to-normal vision and they were provided with a subsistence allowance. The study was undertaken with the understanding and informed written consent of each participant and the experimental procedures were approved by the Institutional Review Board of New York University Abu Dhabi.

As there are no previous studies investigating Weber’s law in a two-finger pointing movement upon which to refer for an a priori power analysis, the sample size was based on similar studies investigating Weber’s law in action tasks (e.g., Bruno et al., 2016; Derzsi & Volcic, 2023; Franz et al., 2009; Ganel et al., 2008, 2014; Utz et al., 2015). To estimate the final inter-finger separation variability more precisely, we have substantially increased the number of trials per participant.

### Apparatus

Participants were tested individually in a dimly-lit room, and were seated on a height adjustable chair so that the thorax pressed gently against the front edge of the table (75 × 120 cm, Thorlabs Inc.). Stimulus presentation was controlled by PsychToolbox-3 (Brainard & Vision, 1997; Kleiner et al., 2007; Pelli, 1997). The targets were presented on a touchscreen (ASUS, 15.6” FHD IPS display, 1920 × 1080, 60 Hz) lying on and firmly fixed to the table (see Figure 1A). The center of the touchscreen was positioned 16 cm away from the lower edge of the table, and 33 cm from a starting point (a 6 mm high screw with a diameter of 10 mm). Targets consisted of two circular dots (10 mm diameter) or two circular dots connected by a line (1.5 mm in thickness), placed at different distances from each other (40, 55, 70, 85, 100 mm). Targets were aligned along the *y* axis, and their center point corresponded with the center of the touchscreen. Targets’ position along the *x* axis shifted in the range of ±10 mm to avoid participants always reaching the same locations in space. Participants’ vision was controlled by means of liquid-crystal occlusion goggles (Red Scientific, Salt Lake City, UT, USA).

To track the kinematics of the participants’ right hand, we used the Optotrak Certus system (Northern Digital Inc., Waterloo, Ontario, Canada) controlled by the MOTOM toolbox (Derzsi & Volcic, 2018). Data were acquired online at 100 Hz, with an accuracy of 0.1 mm and resolution of 0.01 mm. Three infrared-emitting diodes were attached to the following points on participants’ right upper limb: (1) wrist (radial aspect of the distal styloid process of the radius), (2) thumb (ulnar side of the nail), and (3) index finger (radial side of the nail).

### Procedure

Before each trial, participants held the ulnar side of the hand on the table, with the index finger and thumb pinched together on the starting point. As soon as two dots (Points condition) or two dots connected by a line (Line condition) appeared, participants were asked to reach out and simultaneously touch the dots with the index finger and thumb, as fast and accurately as possible. Once participants started the movement and the hand reached a distance of 5 cm (along the *x*-axis) from the starting point, the occlusion goggles changed from a transparent to an opaque state, and participants had to conclude the action without vision (see Figure 1A). Once the movement was completed, participants were required to put their hand back on the starting point, and only then the occlusion goggles became transparent again.

Each condition consisted of 240 trials (48 trials per inter-target distance). The two conditions were separated by a break of at least 5 minutes. The order of conditions was counterbalanced across participants. The experimental session (preceded by 20 practice mixed trials) lasted approximately 60 minutes.

### Data Analysis

Data analysis was performed using R (R Core Team, 2024). The raw data were smoothed and differentiated with a third-order Savitzky-Golay algorithm (window size = 11 points), to reconstruct the 3-D positions, velocities and accelerations of each marker as a function of time. Movement onset was calculated as the time of the lowest, non-repeating wrist acceleration value prior to the continuously increasing wrist acceleration values (Volcic & Domini, 2016). The end of the movement was defined as the time at which the index and the thumb finger made contact with the targets, and quantified on the basis of the Multiple Sources of Information method (Schot et al., 2010). In particular, we considered the following criteria: the 3D velocities of the index finger and the thumb, and their velocities and positions on the *x* and *z* axes. Moreover, the probability of a moment being the end of the movement decreased over time to capture the first instance in which the above criteria were met. The final inter-finger separation was computed as the Euclidean distance between the index and the thumb at the end of the movement.

Trials in which the end of the movement was not captured correctly were discarded from analysis. The exclusion of these trials (223 trials, 2.3% in total) left us with 9377 trials for the final analysis.

We focused our analyses on the average final inter-finger separation and its SD, the latter being the key measure in the present study. Data were analyzed using Bayesian distributional linear mixed-effects models. Unlike conventional regression models assuming a constant SD across observations, a distributional regression model estimates the location parameters, as well as their SD. The model included as fixed-effects the categorical variable Condition (Points vs. Line) in combination with the continuous variable Inter-target distance, which was centered before being entered into the model. The model included independent random effects for subjects to capture the dependency among data due to the repeated measures design. The model was fitted considering weakly informative prior distributions for each parameter. Four Markov chains were simultaneously run, each for 4,000 iterations (1,000 warm-up samples), for a total of 12,000 post-warm-up samples.

The obtained posterior distributions quantify the uncertainty about each estimated parameter (intercepts and slopes) conditional on the priors, model, and data. We have summarized them by computing their means and their 95% credible intervals (95% CrI). To compare estimates between two conditions, we have computed the credible difference distributions by subtracting their respective posterior distributions, which were again summarized by computing their means and their CrI. For statistical inferences we assessed the overlap of the 95% CrI with zero. Chain convergence was assessed using the 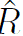 statistic (all values equal to 1) and visual inspection of the chain traces. To test the predictive accuracy of the fitted models, we performed a leave-one-out cross-validation by using the Pareto Smoothed Importance Sampling. All Pareto k values were below 0.5.

## Results

Figure 2A represents the final inter-finger separation as a function of the inter-target distance. These raw data and their fit show that in the Points condition, the final inter-finger separation scaled with the inter-target distance (slope = 0.806, 95% CrI [0.742, 0.870]; intercept = 70.582, 95% CrI [64.847, 76.109]). The Line condition exhibited a similar pattern (slope = 0.819, 95% CrI [0.763, 0.876], intercept = 71.382, 95% CrI [66.697, 75.995]). No credible difference between the Point and Line conditions was found neither in terms of slopes (0.013, 95% CrI [−0.048, 0.073]) nor intercepts (0.799, 95% CrI [−2.991, 4.713]).

**Figure 2.**
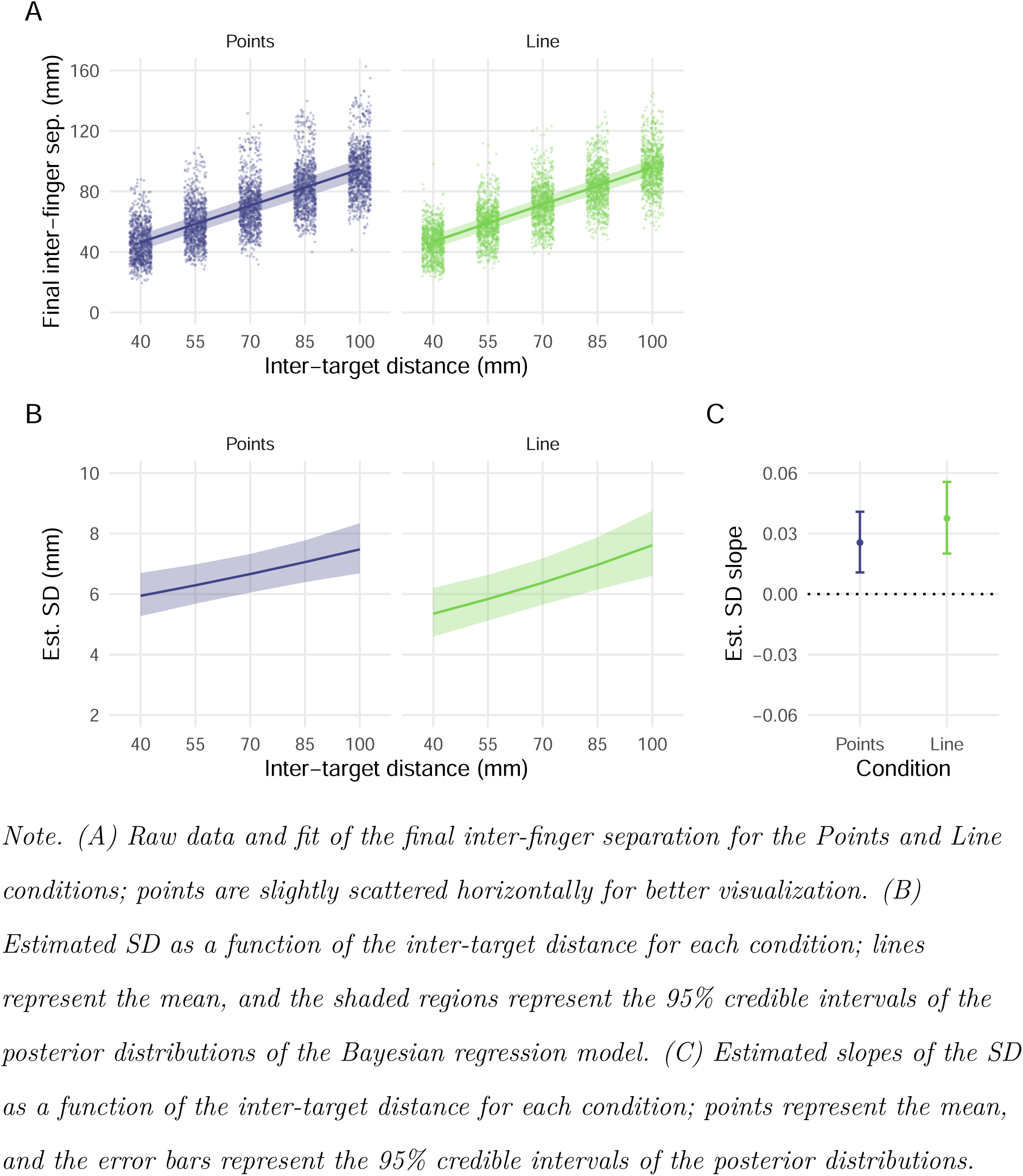
Experiment 1 Results.

Crucially, the SD of the final inter-finger separation (Figure 2B) increased as a function of the inter-target distance both in the Points (slope = 0.026, 95% CrI [0.011, 0.041]; intercept = 6.662, 95% CrI [6.053, 7.322]) and in the Line condition (slope = 0.038, 95% CrI [0.020, 0.056]; intercept = 6.375, 95% CrI [5.652, 7.176]). The estimated slopes for the Points and Line conditions did not differ from each other (0.012, 95% CrI [−0.010, 0.034]; and they showed positive and substantially above zero 95% CrIs (Figure 2C). The two conditions did not differ even in terms of intercepts (−0.286, 95% CrI [−1.111, 0.575])

In summary, our results indicate that two-finger pointing actions are not immune to Weber’s law, in sharp contrast with the predictions of the TVS and DIS hypotheses. The additional prediction that the stimulus type might influence the emergence of Weber’s law was not substantiated. The Points and Line conditions yielded comparable results.

## Experiment 2

The SMC hypothesis holds that performing a movement with vision available throughout the action (closed-loop) should eliminate, or at least reduce, the effects of Weber’s law, because available feedback would promote online adjustments and support motor calibration and, thus, affect movement variability. Instead, the availability of vision should not affect the predictions of the other hypotheses (see Figure 1B, Exp. 2). Therefore, to assess the role of visual feedback, in Experiment 2, we replicated Experiment 1 with the single change that visual input was now consistently available.

## Methods and Materials

All methodological aspects were the same as in Experiment 1 except for what follows.

### Participants

Twenty participants (mean age = 20.1 years, SD = 1.5 years, eight males) were recruited for this experiment. None of them had participated in Experiment 1.

### Procedure and Data Analysis

Occlusion goggles were not used, thus, participants had full vision of the targets and their own hand before, during and after each action. A total of 345 trials (3.6% in total) were discarded from the analysis, which left us with 9255 trials.

## Results

Figure 3A represents the final inter-finger separation as a function of the inter-target distance. These raw data and their fit show that in the Points condition, the final inter-finger separation scales with the inter-target distance (slope = 0.922, 95% CrI [0.890, 0.954]; intercept = 76.166, 95% CrI [74.222, 78.112]). The Line condition exhibits a similar pattern (slope = 0.928, 95% CrI [0.905, 0.952]; intercept = 75.964, 95% CrI [74.196, 77.776]). No credible difference between the Point and Line conditions was found neither in terms of slopes (0.006, 95% CrI [−0.020, 0.032]) nor intercepts (−0.202, 95% CrI [−1.471, 1.084]).

**Figure 3.**
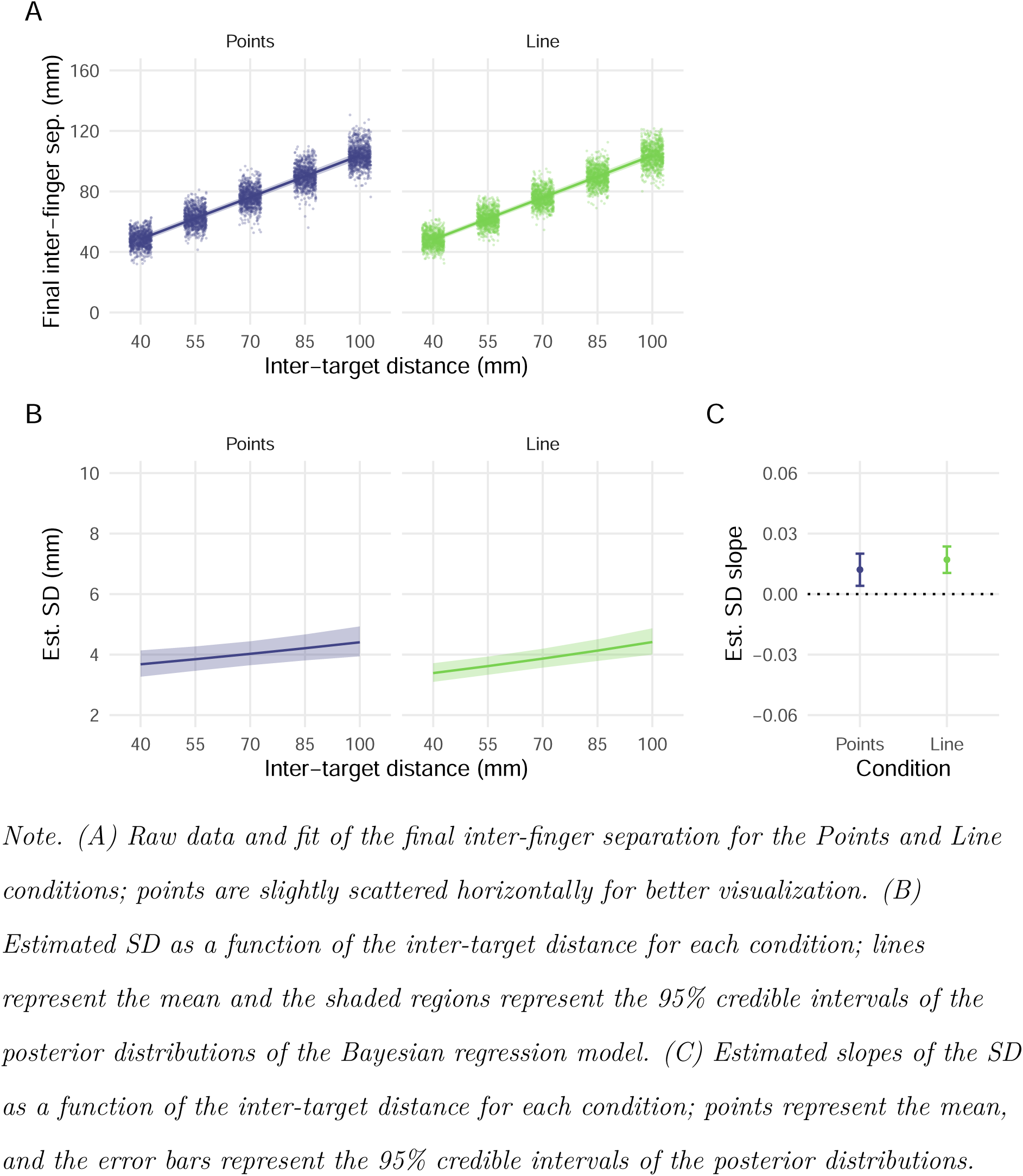
Experiment 2 Results.

Crucially, the SD of the final inter-finger separation (Figure 3B) increased as a function of the inter-target distance both in the Points (slope = 0.012, 95% CrI [0.004, 0.020]; intercept = 4.025, 95% CrI [3.651, 4.438]) and in the Line condition (slope = 0.017, 95% CrI [0.010, 0.024]; intercept = 3.866, 95% CrI [3.569, 4.193]). The estimated slopes for the Points and Line conditions did not differ from each other (0.005, 95% CrI [−0.005, 0.015]) and they showed positive and substantially above zero 95% CrIs (Figure 3C). The two conditions did not differ even in terms of intercepts (−0.158, 95% CrI [−0.571, 0.237]).

In summary, these results replicate those observed in Experiment 1 and are again in contrast with the predictions of the TVS and DIS hypotheses. Weber’s law was found in both Points and Line conditions even when vision was constantly available. In comparison with Experiment 1, the SD of the final inter-finger separation was substantially reduced and its increase as a function of distance was less steep. These findings partially support the SMC hypothesis by providing evidence of the significant role that available visual feedback plays. However, this feedback alone does not seem sufficient to completely eliminate the effect of Weber’s law.

## Discussion

The main aim of this study was to contrast the existing hypotheses explaining the violation of Weber’s law in grasping by introducing a novel action task (i.e., the two-finger pointing task) that entails the same effectors as a grasping task (i.e., thumb and index finger), but it offers the advantage of measuring precision at the exact moment the goal of the action is attained. Here we found that actions obey Weber’s law. The SD of the final inter-finger separation increased as a function of the inter-target distance, with a slope ranging from 0.012 to 0.038, in line with the classic and modern literature on Weber’s law (e.g., Bhatia et al., 2022; Derzsi & Volcic, 2023; Fechner, 1860; Hesse et al., 2021; McKee & Welch, 1992; Teghtsoonian, 1971).

The violation of Weber’s law in action was proposed as compelling evidence supporting the TVS (Ganel, 2015; Ganel et al., 2008, 2014; Goodale & Milner, 1992; Goodale et al., 2004, 2005). However, our results cast doubt on this interpretation, because Weber’s law is regularly at play when considering a measure closely related to the goal of the action (i.e., the final inter-finger separation). In prior studies, the violation of Weber’s law in action has mainly been examined in grasping tasks by measuring the SD_MGA_. Given that we employed a two-finger pointing task, we cannot rule out the possibility that only the shaping of grip aperture relies on absolute metrics. However, it should be noted that recent studies have suggested that SD_MGA_ may not accurately reflect the precision with which visual size is discriminated. In particular, Bhatia et al. (2022) demonstrated that using MGA as a proxy for the corresponding final estimation response can erroneously mask Weber’s law because the response function of MGA is non-linear. When this non-linear scaling is appropriately considered, Weber’s law can be detected even in grasping movements. Moreover, when other variables than MGA are considered, Weber’s law is regularly observed (e.g., Derzsi & Volcic, 2023; Hesse et al., 2021). For instance, Derzsi and Volcic (2023) found that to achieve stable grasps on differently sized objects the SD of grasp contact positions increases as a function of object size, coherently with Weber’s law. As a final caveat, we must add that MGA, unlike the final inter-finger separation, is not a real final estimation response, and its variability could be strongly influenced by a mix of sensory and motor noise that could cancel out the effects of Weber’s law.

Our findings are also in contrast with the DIS hypothesis (Smeets & Brenner, 1999, 2008), which would have predicted a violation of Weber’s law. However, we should note that the DIS hypothesis does not exclude that, under certain circumstances, even non-positional information (such as size or distance) can be involved in movement processing (Smeets et al., 2020). For two-finger pointing movements, non-positional information (i.e., the inter-target distance) exerts its influence, but further research is needed to identify the specific processes underlying this empirical observation.

The present results are in line with the idea that a common visual representation, i.e., a shared underlying representation of visual information, informs both perception and action (Franz, 2001; Franz & Gegenfurtner, 2008) and other factors can occasionally lead to a dissociation (e.g., BF hypothesis). Once these factors are removed, the effects that are considered unique to perceptual tasks also appear in tasks that involve actions (Hesse et al., 2016; Kopiske et al., 2016).

The SMC hypothesis is, to some extent, also consistent with our results. Specifically, we observed a stronger adherence to Weber’s law when visual feedback was not available (Experiment 1). For this reason, as suggested by previous literature, controlling available feedback represents a necessary step to investigate Weber’s law in action (Bruno et al., 2016; Hesse & Franz, 2009; Schenk & Hesse, 2018). Interestingly, in Experiment 2, Weber’s law was not completely eliminated, despite participants had access to visual feedback before, during and after the movement. A possible explanation for this result may lie in the randomized positioning of targets, which could have slowed down motor calibration.

One potential limitation of our study may stem from the use of digitally presented stimuli. Experimental evidence suggests that grasping 2D digital objects could induce atypical behavior, diverging from natural grasping kinematics (Ozana & Ganel, 2019a, 2019b; Ozana et al., 2020; Westwood et al., 2002). Indeed, the lack of depth and realism makes 2D digital objects impossible to be grasped or manipulated in the same manner as three-dimensional objects. Consequently, 2D digital objects may not be suitable stimuli for studying grasping kinematics. If this is undisputed for grasping, it is not necessarily the case for pointing movements. Indeed, pointing movements do not necessarily require a physical 3D object for realistic execution. In everyday life, pointing is directed toward a wide range of targets, whether in 2D or 3D, such as buttons, switches, and digital elements on screens or tablets. Therefore, the use of digital targets to investigate pointing movements represents a practical and ecologically valid approach, as highlighted by their application in many motor control studies (e.g., Connolly et al., 2003; Diedrichsen et al., 2004; Gentilucci et al., 1996; Hesse et al., 2012, 2014; Pélisson et al., 1986; Rossit et al., 2018).

In conclusion, future research should consider tasks in which Weber’s law is assessed at action completion. This approach would offer deeper insights into the intricate interplay between perception and action and contribute to this long-standing debate.

## Data Availability

All data and R code are publicly available in the Open Science Framework repository (osf.io/w5ujf/).

## Acknowledgments

We acknowledge the support of the NYU Abu Dhabi Research Enhancement Fund (grant RE183), the NYUAD Center for Artificial Intelligence and Robotics, funded by Tamkeen under the NYUAD Research Institute Award CG010, and, the NYUAD Center for Brain and Health, funded by Tamkeen under the NYUAD Research Institute Award CG012.

